# A scalable permutation approach reveals replication and preservation patterns of gene coexpression modules

**DOI:** 10.1101/029553

**Authors:** Scott C. Ritchie, Liam G. Fearnley, Gad Abraham, Michael Inouye

## Abstract

Gene coexpression network modules provide a framework for identifying shared biological functions. Analysis of topological preservation of modules across datasets is important for assessing reproducibility, and can reveal common function between tissues, cell types, and species. Although module preservation statistics have been developed, heuristics have been required for significance testing. However, the scale of current and future analyses requires accurate and unbiased p-values, particularly to address the challenge of multiple testing. Here, we developed a rapid and efficient approach (*NetRep*) for assessing module preservation and show that module preservation statistics are typically non-normal, necessitating a permutation approach. Quantification of module preservation across brain, liver, adipose, and muscle tissues in a BxH mouse cross revealed complex patterns of multi-tissue preservation with 52% of modules showing unambiguous preservation in one or more tissues and 25% showing preservation in all four tissues. Phenotype association analysis uncovered a liver-derived gene module which harboured housekeeping genes and which also displayed adipose and muscle tissue specific association with body weight. Taken together, our study presents a rapid unbiased approach for testing preservation of gene network topology, thus enabling rigorous assessment of potentially conserved function and phenotype association analysis.

## Introduction

Modern high-throughput technologies generate a large amount of data, including genomic, transcriptomic, metabolomic, and proteomic measurements. Rather than consider each component in isolation, network inference techniques are an increasingly common and useful way to integrate omic data to identify meaningful biological relationships within and between molecular systems. These approaches are widely applied to various biological problems, including identification and characterization of gene modules, gene regulatory networks, protein-protein interactions, and prediction of diverse molecular interactions (Abraham et al., 2014; Barabási et al., 2011; Lusis and Weiss, 2010; Schadt, 2009). Typically, one is interested in investigating particularly interesting subsets of these inferred networks, referred to as network *modules* (Gustafsson et al., 2014; Rotival and Petretto, 2014). In this area, analysis of gene coexpression network modules is a particularly useful and common approach (Chen et al., 2008; Emilsson et al., 2008; Fuller et al., 2007; Inouye et al., 2010a, 2010b; Keller et al., 2008; Ritchie et al., 2015; Yang et al., 2009; Zhang et al., 2013).

Of particular interest are studies that have examined the preservation of gene coexpression network modules across datasets. Module preservation analysis has enabled researchers to assess module reproducibility (Emilsson et al., 2008; Fuller et al., 2007; Hawrylycz et al., 2012; Miller et al., 2010; Xia et al., 2006), to determine how modules change across conditions (Fuller et al., 2007; Keller et al., 2008; Van Nas et al., 2009), to examine module tissue specificity (Cai et al., 2010; Keller et al., 2008), and to identify modules conserved across different species (Boyle et al., 2014; Gerstein et al., 2014; Stuart et al., 2003). The need for scalable module preservation analysis is rapidly increasing as large multi-omic datasets with dozens of tissues, cell-lines, and corresponding metadata become common and openly available. For example, module preservation analysis has recently been applied to pilot data from the Genotype Tissue Expression (GTEx) project (The GTEx Consortium, 2013) with the aim of identifying tissue-specific and cross-tissue gene coexpression modules (Mele et al., 2015; Pierson et al., 2015; The GTEx Consortium, 2015).

Although there have been intense efforts to design and improve methods used for gene network inference (Marbach et al., 2012; Prill et al., 2010), rigorous statistical methodology for assessing preservation of the subsequent gene modules of interest has received far less attention. The preservation of gene coexpression modules is typically quantified via visual inspection and/or tabulation of the module genes in an independent dataset (Boyle et al., 2014; Gerstein et al., 2014; Keller et al., 2008; Miller et al., 2010; Van Nas et al., 2009; Xia et al., 2006). Although this type of approach can be informative, it cannot capture information about network topology, *i.e.* the relationships between genes, thus it potentially discards important biological information. For example, genes that are highly connected are often essential to an organism’s survival (Carlson et al., 2006; Jeong et al., 2001), and node connectivity is associated with relative biological importance (Horvath and Dong, 2008; Langfelder et al., 2013).

Here, we are primarily concerned with approaches based on weighted gene coexpression network analysis (Zhang and Horvath, 2005). Here, network inference includes two key matrices: a coexpression matrix and an adjacency matrix. For a dataset with *n* genes assayed for *m* samples, the coexpression matrix is an *n × n* matrix containing the correlation coefficient between each pair of genes. The adjacency matrix (also *n × n*) is a transform of the coexpression matrix that contains values between 0 and 1, denoting the connection strength between each pair of genes. A power transform is a typical approach for calculating this adjacency matrix, as exponentiating each entry of the coexpression matrix penalises relatively weak correlations toward zero. Gene module detection is typically performed on the adjacency matrix using a clustering algorithm that identifies tightly connected gene modules, which may represent biological pathways. While common tools such as WGCNA (Langfelder and Horvath, 2008) generally follow the above methodology, it is important to note that preservation statistics do not assume a particular approach to network inference or module definition.

A recent study by Langfelder *et al* developed a suite of statistics for quantifying replication and preservation of gene coexpression module topology (Langfelder et al., 2011). For convenience, definitions of the Langfelder and Horvath statistics are given in the **Experimental Procedures** and their biological interpretation in the **Supplemental Materials**. Broadly speaking, these statistics measure the preservation of module *density* and *connectivity*. *Density* statistics assess whether a module is highly connected and has coherent expression in the test dataset. *Connectivity* statistics assess whether the gene-gene relationships (*i.e.*, coexpression, connectivity, module membership of each gene) are preserved between the discovery and test datasets (Langfelder et al., 2011).

The current approach for assessing statistical significance of module replication and preservation assumes normality of the statistics under the null, and relies on heuristic significance thresholds (Langfelder et al., 2011). However, the increasing scale of current module preservation analyses requires both rapid and statistically rigorous methods to overcome substantial barriers in differentiating signal from noise. In particular, studies are now performing unbiased discovery of preserved gene coexpression modules, thus incurring a large multiple hypothesis testing penalty (Cai et al., 2010; Pierson et al., 2015)., It is crucial that preservation p-values are unbiased and accurately calibrated in order to control type I and type II error rates and accommodate multiple-testing adjustment (Bender and Lange, 2001).

Permutation testing is a commonly used approach to assessing statistical significance in the absence of distributional assumptions, and involves empirical generation of null distributions by random sampling of gene sets in the test dataset. However, at least *w* permutations are required to estimate significance at a threshold of *1/w* (Phipson and Smyth, 2010). The analysis of large datasets, together with the concomitant multiple testing correction burden, necessitates increasingly stringent significance thresholds, making permutation-based significance testing computationally challenging.

Here, we address this challenge by developing a rapid and efficient approach for assessing module preservation, available as an R package *NetRep*. We use *NetRep* to create and assess the empirical null distributions of Langfelder *et al.*’s suite of module preservation statistics. We show the majority of these statistics are non-normal and thus in need of a permutation approach. Next, we demonstrate *NetRep*’s scalability to large-scale module preservation analysis by analysing a publicly available resource of mouse adipose, brain, liver, and muscle tissue expression (Yang et al., 2006). We quantify cross-tissue gene coexpression module preservation to identify and characterize multi-tissue modules associated with mouse body weight. Consequently, we uncover a body weight-associated module with differential adipose and muscle tissue expression.

## Results

### Rapid module preservation analysis

We have developed a time and memory efficient method for massively parallel calculation of module preservation statistics. The software is available as an R package, *NetRep*, which can be downloaded from https://github.com/InouyeLab/NetRep. Implementation details are provided in the **Supplemental Experimental Procedures**.

To examine the null distributions of the module preservation statistics, we applied *NetRep* to publicly available gene expression data for brain, adipose, liver, and muscle tissues from a BxH mouse cross (Yang et al., 2006). From 334 total mice, there were 249 brain, 295 adipose, 306 liver, and 319 muscle tissue samples available for analysis (**Experimental Procedures**). **Figure 1** illustrates the workflow of network construction, module detection, and module preservation analysis.

**Figure 1:**
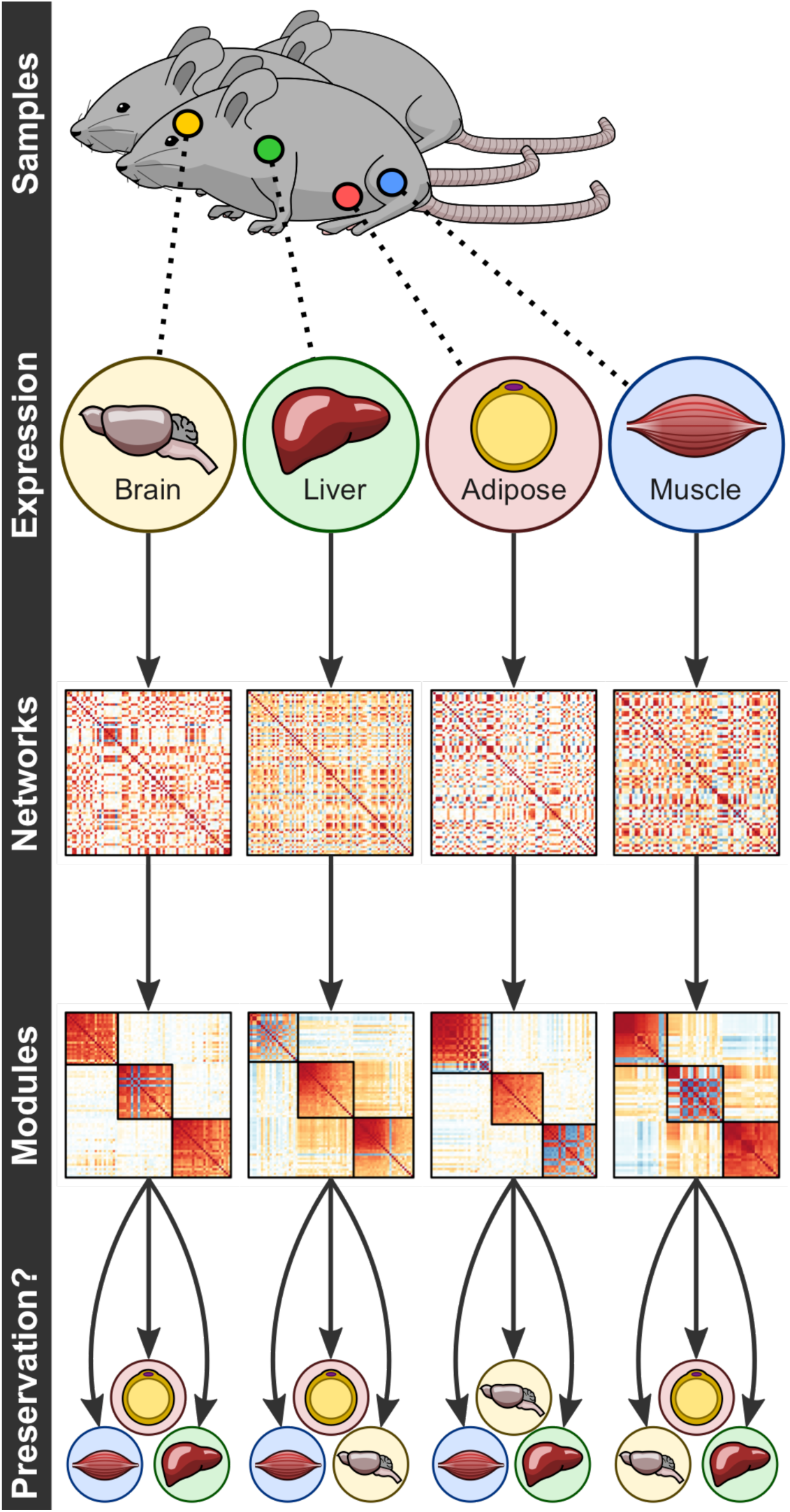
Workflow of network construction, module detection, and module preservation analysis workflow. First, coexpression networks were calculated from the gene expression in each tissue. Next module detection was performed using weighted gene coexpression network analysis (WGCNA) (**Experimental Procedures**). Finally, the preservation of each module was assessed in each non-discovery tissue.

We inferred weighted gene coexpression networks (**Experimental Procedures**, (Zhang and Horvath, 2005)) for each tissue, identifying 38, 66, 29, and 32 distinct modules in the brain, liver, adipose, and muscle tissues, respectively (**Figure S1**). Coexpression networks were calculated for each tissue using the pairwise Pearson correlation coefficient. Adjacency networks were defined as the absolute value of the correlation coefficient exponentiated to powers 12, 5, 4, and 5 respectively, selected through the scale-free topology criterion (**Experimental Procedures**, (Zhang and Horvath, 2005)) in each respective tissue (**Figure S2**). Runtime comparison of *NetRep* versus WGCNA’s *modulePreservation* function for these modules is provided in the **Supplemental Experimental Procedures** (see also **Figure S3**).

### Null distributions of module preservation statistics

We investigated the normality of the seven module preservation statistics (**Experimental Procedures**) by comparing permutation-based null distribution quantiles to theoretical normal distribution quantiles (**Figure S4**). For each discovered module, module preservation statistics were calculated on 100,000 random gene sets of identical size in each non-discovery tissue. Across all 38 brain modules, 66 liver modules, 29 adipose modules, and 32 muscle modules, 495 null distributions were generated for each module preservation statistic. We observed strong non-normality of null distributions generated for the *module density*, *mean coexpression*, and *correlation of coexpression* statistics (**Figure S4**). Moderate non-normality was also observed in null distributions for the other four statistics. We also observed increasing non-normality with decreasing module size, particularly for modules of <100 genes (**Figure S4**).

Matched Z-score p-values and permutation p-values were calculated for each module and preservation test statistic. Substantial inflation of Z-score p-values was observed for *module density*, *mean coexpression*, and *correlation of coexpression* (**Figure 2**), consistent with observed non-normality. Moderate inflation also observed for *proportion of variance explained* and *mean module membership* (**Figure 2**). These results are consistent with the extremely low p-values that motivated heuristic significance thresholds (Langfelder et al., 2011).

**Figure 2:**
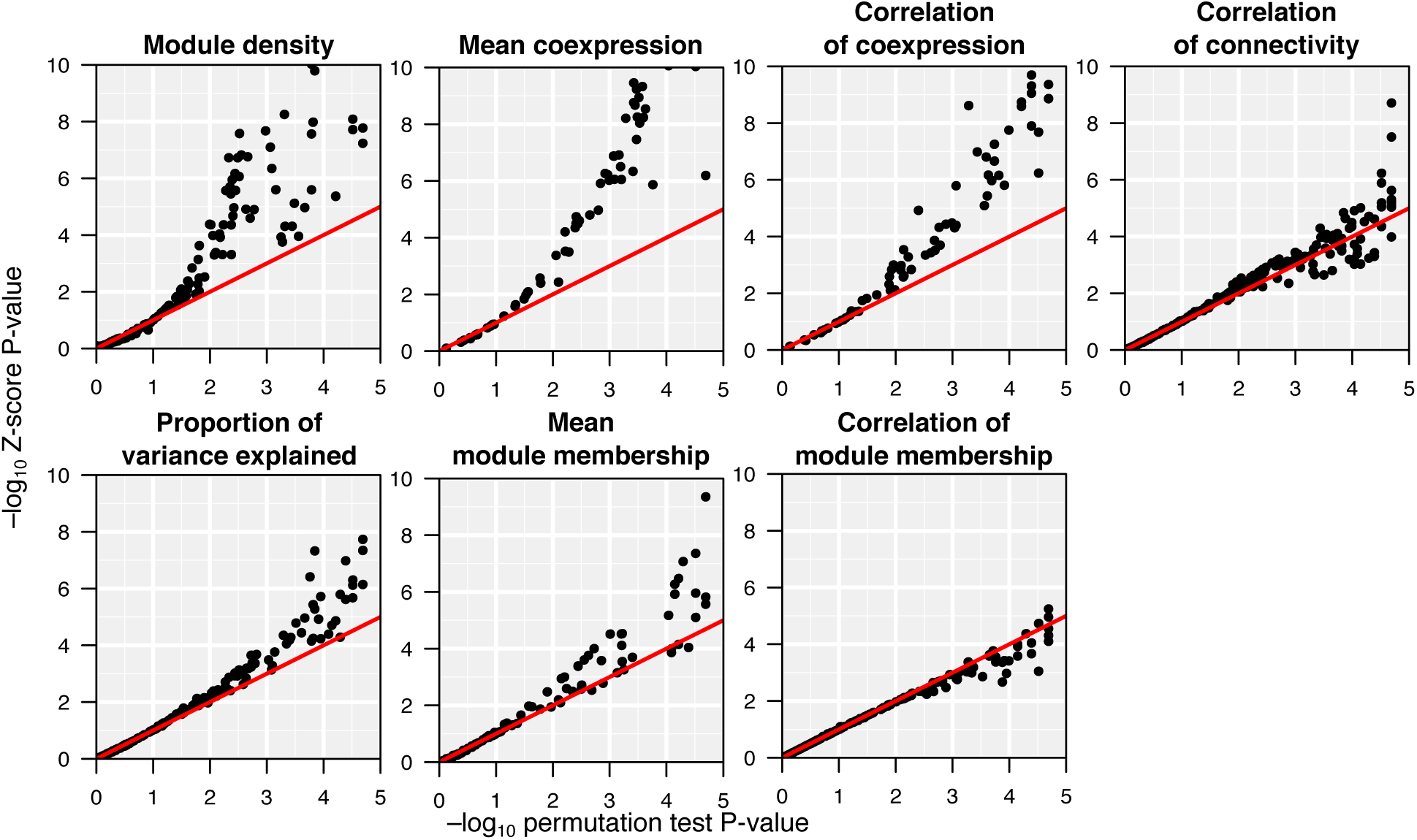
Comparison of permutation test p-values to Z-test p-values for each of the 165 modules when tested in their 3 non-discovery tissues. The mean and standard deviation of the 495 null distributions were used to calculate the Z-test p*-*values. Null distributions were generated from 100,000 permutations. P-values were plotted on a −log10 scale. Coexpression tests with −log10 (*P*_Z-test_) > 10 are not plotted (25 data points for module density, 11 for correlation of coexpression, and 21 for mean coexpression).

Deviation from normality was most extreme for the *module density* statistic, thus we investigated whether the scale-free assumption used to define the adjacency network was a contributing factor (Zhang and Horvath, 2005). If the global connectivity of the adjacency network follows a power law, then it is said to be *scale-free*, thus the assumption is that a few *hub* genes are strongly connected in the network while most genes are only weakly connected (Barabási and Albert, 1999; Stumpf and Porter, 2012). To test this, we generated null distributions from 10,000 permutations in the muscle tissue for the 66 liver modules, varying the soft-threshold power from 12 to 1. We observed a trend towards normality as the soft-threshold power approached 1, indicating that the scale-free assumption contributes to non-normality for this statistic (**Figure S5**). We similarly generated null distributions for the *correlation of connectivity* as it is also calculated from the adjacency matrix, however its deviation from normality was only mild (**Figure S6**).

### Cross-tissue module preservation in mouse transcriptomic data

We next examined the preservation of each discovered module in other tissues by evaluating the permutation test p-values for each of the 495 null distributions (**Experimental Procedures**). We defined strong evidence for a module’s preservation in another tissue as all test statistics achieving *P* < 0.0001, weak evidence if one or more, but not all, test statistics were *P* < 0.0001, and no evidence if no test statistics are *P* < 0.0001. The significance threshold of 0.0001 was chosen to adjust for the 495 tests performed for each preservation statistic. **Figure 3** provides a summary view of the cross-tissue module preservation in the BxH mice and **Figure 4** shows the preservation evidence for each module in each non-discovery tissue.

**Figure 3:**
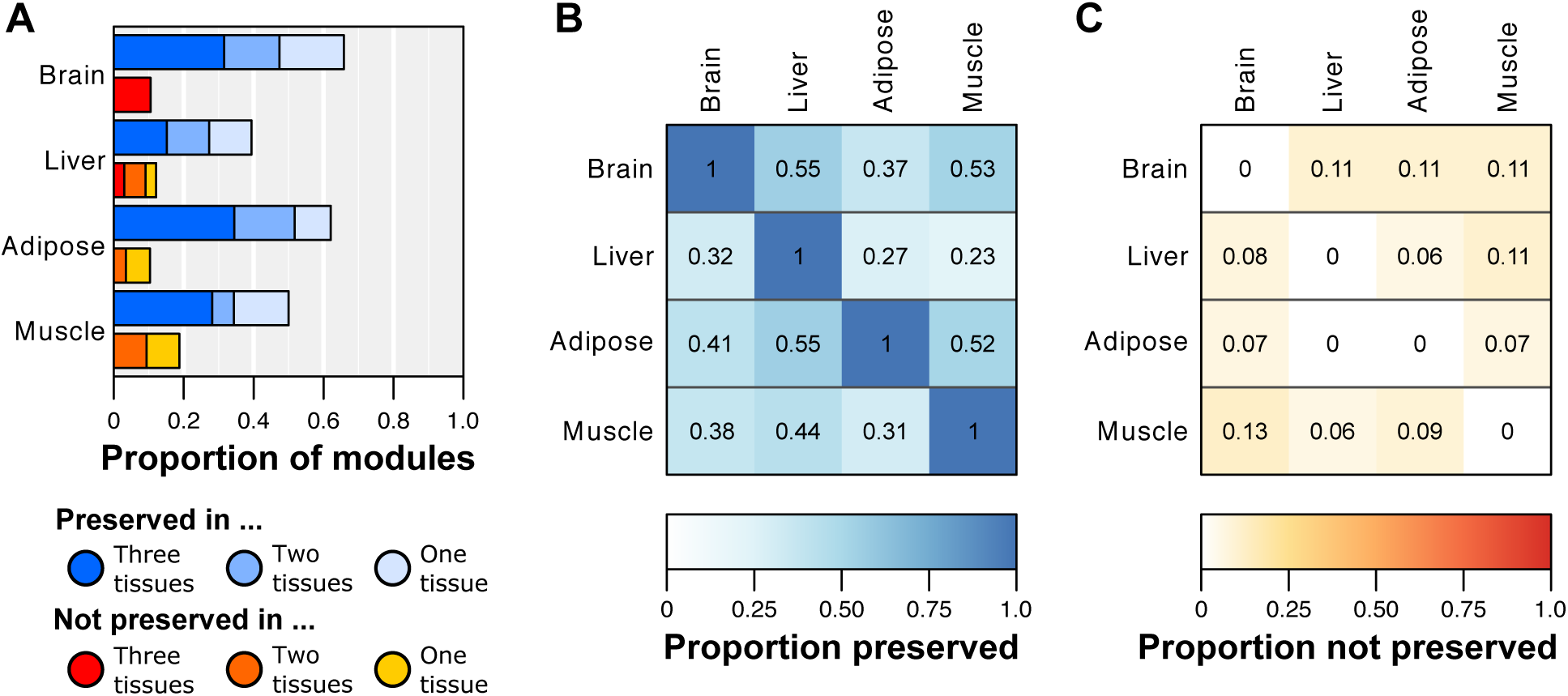
Summary of cross-tissue module preservation. (A) Summary of evidence of preservation for modules discovered in each tissue. Here, a module was considered *preserved* if it had strong evidence of preservation in another tissue, and *not preserved* if it had no evidence of preservation in another tissue. (B) Tissue similarity based on the proportion of modules discovered in the row tissue with strong evidence of preservation in the column tissue. (C) Tissue uniqueness based on the proportion of modules discovered in the row tissue with no evidence of preservation in the column tissue. Note that the heatmap entries in panels B and C cannot be read vertically.

**Figure 4:**
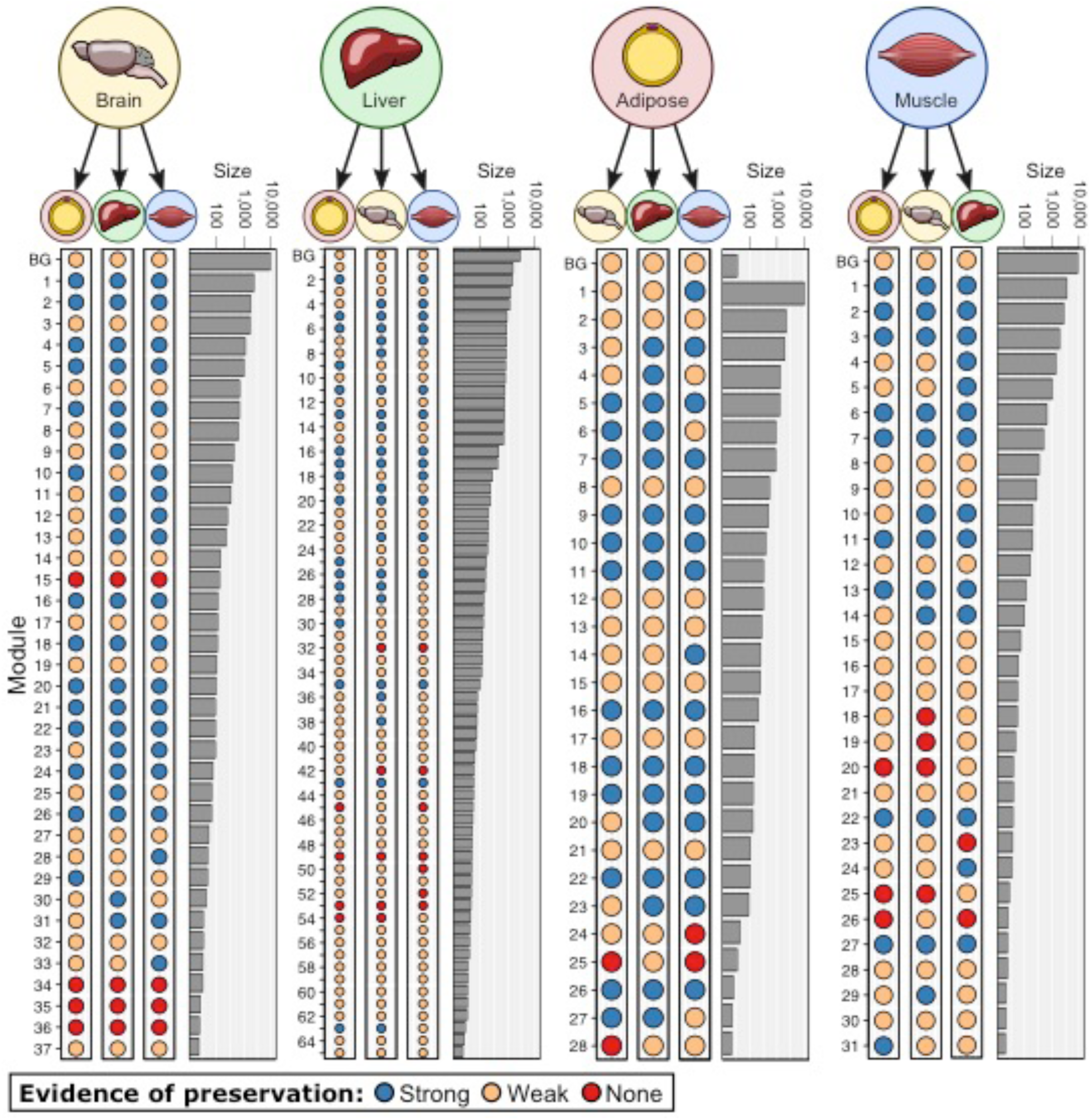
Evidence for preservation for each module in each non-discovery tissue. Module sizes (horizontal bar plots) are shown on a log10 scale. Modules (dots) are sorted according to module size and coloured according to preservation evidence (‘strong’ evidence is blue, ‘weak’ is yellow, ‘none’ is red).

We observed widespread preservation for modules in all four tissues (**Figure 3a,b)**. 85 of 165 modules (52%) had strong evidence of preservation in at least one other non-discovery tissue, and 41 modules (25%) had strong evidence of preservation in all non-discovery tissues (**Figure 3a**). In contrast, only 21 modules (13%) had no evidence for preservation in any other tissue, suggesting tissue specificity of these modules (**Figure 3a,c**). The liver had the lowest proportion of modules preserved in at least one other tissue, however, it had twice as many modules in total than any other tissue. Many of these were small (< 100 genes), and had only weak evidence for preservation (**Figure 4**). Only the brain and liver had any modules with no evidence for preservation in all three non-discovery tissues (**Figure 3a, 4**). These results were broadly consistent with recent results from the GTEx consortium, who observed high similarity between coexpression network modules across nine human tissues (including adipose and muscle tissues) (The GTEx Consortium, 2015).

In comparing *NetRep* results with those obtained through summary Z-score and heuristic thresholds (**Experimental Procedures**, (Langfelder et al., 2011)), the two approaches generally obtained similar levels of evidence for preservation (**Table 1**), however differences in preservation were observed for 120 of the 495 (24%) module preservation tests. In terms of module preservation, 55 (24%) of the modules found to be strongly preserved by heuristics were classified as weakly preserved by *NetRep*. Similarly, 44 (54%) of the modules found to be not preserved by heuristic were classified as weakly preserved by *NetRep*.

**Table 1:**
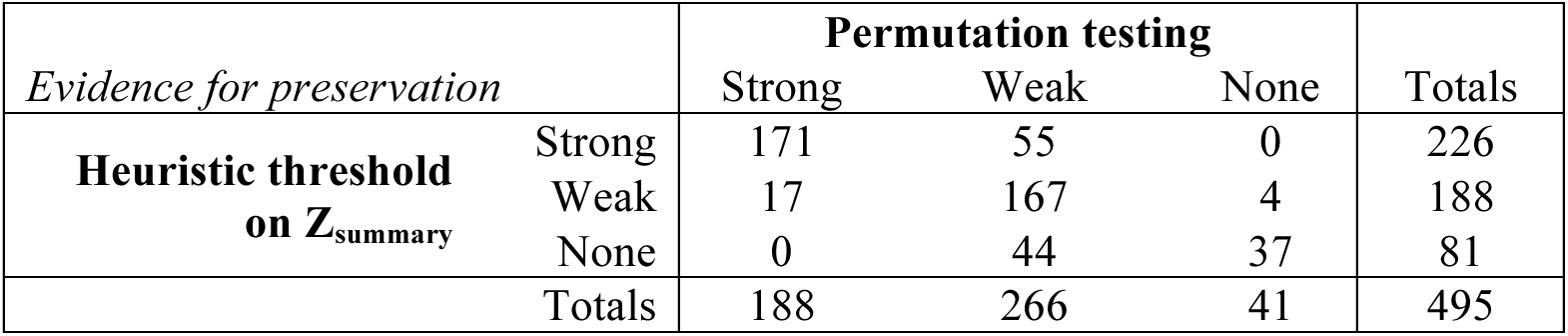
Comparison of module preservation evidence between *NetRep* and heuristic approach.

In total, *NetRep* found that 41 modules (10 discovered in adipose tissue, 12 in brain, 10 in liver, and 9 in muscle) were preserved in all non-discovery tissues. Analysis of Gene Ontology (GO) terms and Kyoto Encyclopedia of Genes and Genomes (KEGG) pathways for each module (**Experimental Procedures**), showed that these were putative housekeeping modules, which were most frequently enriched for genes involved in translation, and to a lesser extent transcription and basic cellular functions, *e.g.*, cell cycle, apoptosis, and DNA repair (**Table S1**). Notably, the putative housekeeping modules were most frequently enriched for genes coding for ribosomal proteins with 10 of 41 putative housekeeping modules enriched for the Ribosome pathway in KEGG (**Table S1**).

### Multi-tissue weight-associated modules

The BxH mice were bred to enhance differentiation of cardiovascular disease risk traits such as obesity and circulating lipids (Yang et al., 2006). Previous coexpression network analysis of the BxH mice focused on the identification of modules associated with mouse weight (Chen et al., 2008; Ghazalpour et al., 2006). We were interested in identifying whether multi-tissue modules, those with strong evidence of preservation in any other tissue, were associated with obesity. We therefore tested each multi-tissue module’s summary expression (1st principal component, **Experimental Procedures**) for an association with mouse weight in any tissue for which there was strong evidence for its preservation (**Experimental Procedures**). Significant association with weight was defined as *P* < 0.0001 (Bonferroni correction). Each regression was adjusted for sex due to its strong effect on both mouse weight and gene expression (Fuller et al., 2007; Ghazalpour et al., 2006; Yang et al., 2006).

Of the 85 multi-tissue modules, 43 modules were significantly associated with mouse body weight in either the discovery tissue or in a tissue where it was strongly preserved, comprising 57 body weight associations in total (**Table S2**). Twenty-seven (32%) of these multi-tissue modules were also putative housekeeping modules. Weight was most frequently associated with modules in adipose tissue (28 of 57 associations) and liver tissue (24 of 57 associations). Notably, there were many cases where multi-tissue modules were not associated with weight in their discovery tissue but displayed association in non-discovery tissues (**Table S2**). In total, 13 multi-tissue modules were associated with mouse weight in multiple tissues (**Table S2**). Interestingly, we observed different directions of weight association across tissues for five modules: *i.e.* in tissue A, an increase in weight was associated with a decrease in module summary expression, but in tissue B an increase in weight was associated with an increase in module summary expression, or vice versa (**Table S2**). For these five modules, visualisation of the network topology indicated the weight-associated differential summary expression reflected differential whole-module expression for two modules: liver module 35 and brain module 20.

We selected liver module 35 (LM35) for further investigation (**Figure 5**). LM35 is a putative housekeeping module consisting of 99 genes (permutation test *P* ≤ 1 *×* 10^−5^ for all statistics in the brain, adipose, and muscle tissues). Consistent with previous analysis of the putative housekeeping modules, GO term and KEGG pathway enrichment indicated LM35 was primarily enriched for ribosomal genes involved in translation (**Table S3**). While a majority of the probes in LM35 lacked annotation for the custom array, 17 of its 24 annotated genes coded for ribosomal proteins (**Table S4** and **Experimental Procedures)**.

**Figure 5:**
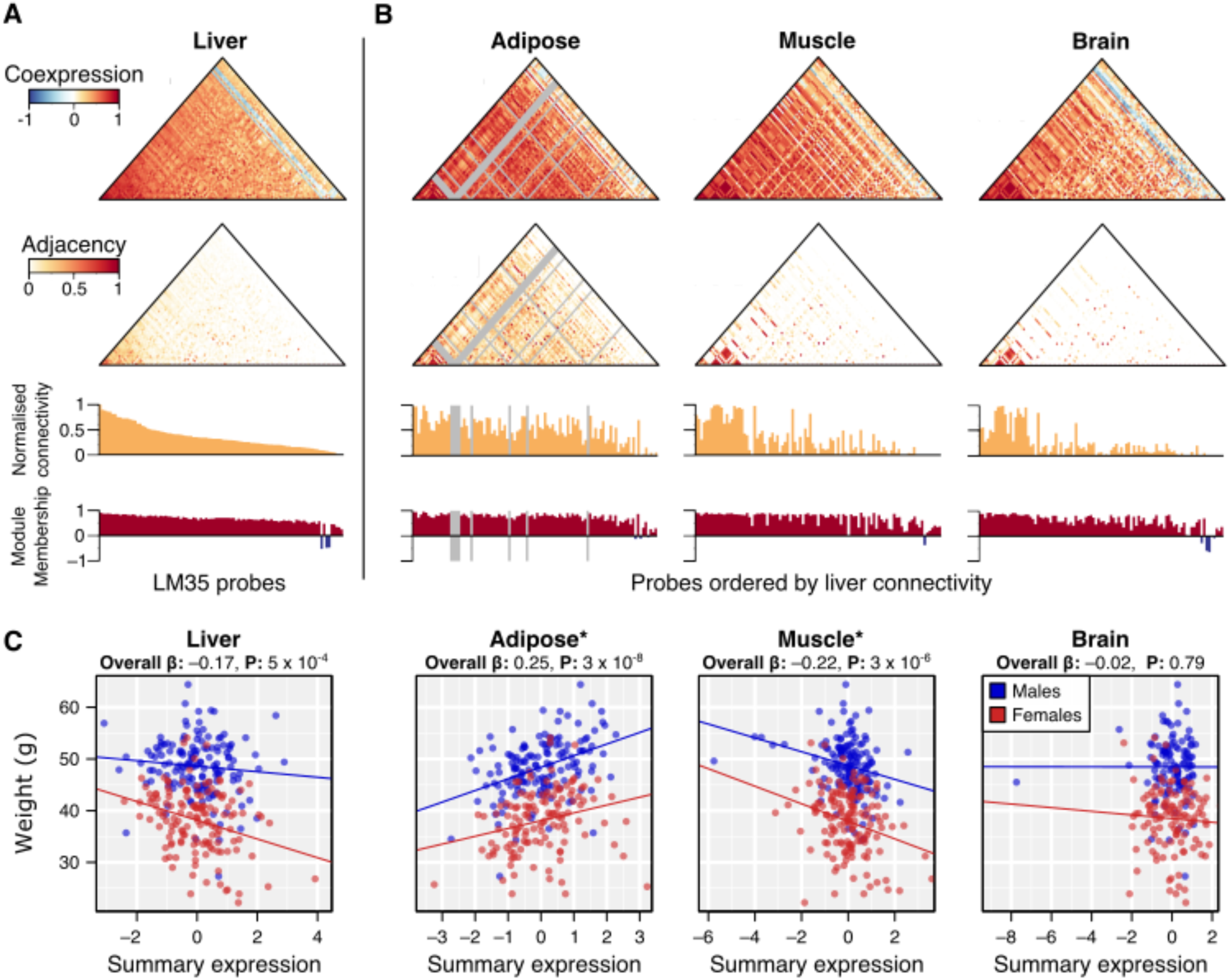
Liver module 35 (LM35). (A) Network topology in the liver (discovery tissue). From top to bottom: heatmap of the pairwise probe coexpression (Pearson correlation), heatmap of the pairwise adjacencies, relative connectivity within each tissue (normalised by maximum value in each tissue), and module membership. Probes are ordered from in descending order of connectivity in liver tissue. (B) Network topology in adipose, muscle, and brain tissues. Probes are ordered as in liver tissue. Grey bars denote missing probes. (C) Scatter plots of standardized LM35 summary expression vs. body weight. Points on the scatter plot are coloured by sex (males in blue, females in red) and linear regression models were adjusted for gender (lines shown are fitted within genders). Models were robust to outliers (summary expression < −3 SD).

Increased body weight was associated with increased LM35 expression in adipose tissue (*P* = 3 *×* 10^−8^) and decreased LM35 expression in muscle tissue (*P* = 3 *×* 10^−6^) (**Figure 5**). Consistent with this, its summary expression profile was negatively correlated between the adipose and muscle tissues (Pearson’s *ρ* = −0.13) and the expression of 64 of its 99 probes were negatively correlated across the two tissues.

We subsequently tested 20 other cardiometabolic traits for association with LM35 expression in either adipose and muscle tissues (**Table S5**). Consistent with the direction of the weight associations, increased insulin, total cholesterol, and total fat were associated with increased adipose expression and decreased muscle expression (FDR *q* < 0.025; **Table S5**). These changes in LM35 expression were also associated with a decrease in the ratio of glucose over insulin (**Table S5**). Increased LM35 adipose expression was associated with increased glucose, other fat, body length, and monocyte chemotactic protein-1 (MCP-1) (**Table S5**). On the other hand, decreased LM35 muscle expression was associated with increased abdominal fat, free fatty acids, total cholesterol, and LDL+VLDL, but a decreased ratio of HDL to LDL+VLDL (**Table S5**).

## Discussion

Accurate and unbiased assessment of the replication and preservation of gene coexpression modules requires permutation testing of network feature similarity (Langfelder et al., 2011). However, the current approach employs heuristics to assess significance due to the computational burden of these calculations (Langfelder et al., 2011). While heuristics may be appropriately employed for a small number of modules, the scale of module preservation and replication analyses now requires a rapid and statistically rigorous method to enable adjustment for multiple hypothesis testing, and consequently allow confident investigation of the underlying biology. In this study, we have developed a rapid and efficient approach for assessing gene coexpression module replication and preservation through permutation testing: *NetRep*. We have empirically shown that module preservation statistics are largely non-normal under the null hypothesis of non-preservation. We next applied *NetRep* to a multi-tissue dataset, observing widespread preservation of gene coexpression modules across four tissues. Housekeeping modules, those preserved in all four tissues, were enriched for genes involved in basic cellular processes, most notably ribosomal genes involved in translation. Subsequent investigation of multi-tissue modules associated with body weight revealed that preserved modules can exhibit differential intra-module expression across tissues, and we have identified a housekeeping module linked to obesity and insulin resistance with increased adipocyte expression and decreased muscle expression in overweight mice.

Previous studies have identified multi-tissue modules driving obesity in mice and humans, with concordant expression across tissues (Chen et al., 2008; Emilsson et al., 2008). Here, we found that multitissue modules may be differentially expressed across tissues with corresponding phenotypic differences. The liver module LM35 exhibited negative, positive and negative associations with body weight in liver, adipose and muscle tissues, respectively. Perhaps consistent with its tissue-specific directions of body weight association, LM35 was enriched for genes encoding ribosomal proteins, which maintain putative housekeeping functions. However, the gene set comprising LM35 was the only multi-tissue housekeeping module which exhibited significant patterns of differential body weight association. Furthermore, LM35 was associated with several obesity related traits, including a decreased ratio of glucose over insulin, suggesting an association with decreased insulin sensitivity. The link between insulin sensitivity, obesity, and adipocytes is well established (Hotamisligil et al., 1993; Kahn and Flier, 2000; Kahn et al., 2006) and, consistent with this link, the adipose expression of LM35 was associated with circulating MCP-1 levels. MCP-1 has been shown to be secreted by adipocyte cells as well as overexpressed in obese mice, and it has been shown to decrease insulin-stimulated glucose uptake *in vitro* (Kanda et al., 2006; Sartipy and Loskutoff, 2003).

Protein–protein interaction networks have shown that tissue-specific proteins tend to interact with universally expressed housekeeping proteins, suggesting their recruitment and modification for tissue-specific function (Bossi and Lehner, 2009). The differential intra-tissue functionality for LM35 may be explained in a similar fashion, *i.e.*, that LM35 strongly coexpresses with a number of obesity-linked genes in adipose and muscle tissue that do not coexpress in the liver tissue where the module was discovered. Multiple plausible scenarios may lead to differential inter-tissue module expression. Module expression in one tissue might cause secretion of signalling molecules that regulate the same module in another tissue, or module expression may respond to changes in circulating metabolites or signalling molecules in different ways across tissues. Genetic variants with tissue-specific effects may also have a role in determining module regulation and phenotypic relationships.

In recent years, studies have begun generating and analyzing datasets containing dozens of tissues and cell types, for example the Genotype-Tissue Expression (GTEx) Consortium (The GTEx Consortium, 2015), and the Immunological Genome (ImmGen) (Shay and Kang, 2013) and Immune Variation (ImmVar) projects (De Jager et al., 2015). Already, multiple module preservation analyses have been performed on the GTEx pilot data (Mele et al., 2015; Pierson et al., 2015; The GTEx Consortium, 2015). With large-scale expression studies increasing in scale and complexity, there is an urgent need for powerful and accurate statistical methodologies which quantify module replication and preservation. Here, we have presented an approach for rapid assessment of gene coexpression module preservation and reproducibility which makes possible unbiased large-scale comparative analysis.

## Experimental procedures

### BxH Mouse Cross

The BxH mouse cross is a publicly available dataset comprising samples from 334 mice bred on an Apolipoprotein E null background in order to enhance the differentiation of cardiovascular disease traits. Data collection protocols are extensively described in (Yang et al., 2006). Briefly, mice were fed on a high-fat, high-cholesterol diet from 8 weeks of age for 16 weeks and sacrificed at 24 weeks of age after fasting for 4 hours. Gonadal adipose (epididymal fat pad in males, perimetrial fat pad in females), whole brain, liver, and skeletal hamstring muscle were collected and immediately frozen in liquid nitrogen. Data were downloaded from Sage BioNetworks at https://www.synapse.org/#!Synapse:syn4497, but are also available through GEO through the following identifiers. Brain: GSE3087, liver: GSE2814, adipose: GSE3086, and muscle: GSE3088.

### Transcriptomic profiling

Tissue samples were homogenised and RNA was extracted, prepared, and hybridised on a custom Agilent array as previously described (Schadt et al., 2003, 2008; Yang et al., 2006). Individual transcript intensities were corrected for experimental variation and normalised within each tissue, then reported as the mean log10 ratio of each individual experiment relative to a pool of RNA comprised of equal aliquots of RNA from the respective tissues of 150 randomly selected mice (He et al., 2003; Yang et al., 2006).

Gene expression microarray probes were subsequently quality controlled on a per-tissue basis. First, probes with more than 5% missingness were excluded, followed by samples with >5% probe-level missingness. Remaining missing information was imputed use a *K*-nearest neighbours algorithm using the R package *impute* (version 1.38.1). 22,808 probes and 295 samples from the adipose tissue, 22,950 probes and 249 samples from the brain tissue, 22,863 probes and 306 samples from the liver tissue, and 22,999 probes and 319 samples in the muscle tissue passed quality control.

In total, 20,367 probes for 18,787 genes corresponded to genes from the NCBI build 36/mm8 annotation release. 539 of the genes have been subsequently withdrawn, as they were not predicted in a later annotation. 3,205 probes on the array corresponded to UniGene clusters lacking annotation (*i.e.*, no corresponding Entrez Identifier) (Schadt et al., 2008).

### Phenotypic profiling

Extensive normalised physiological measurements and metabolic measurements from plasma were provided. Measurements with > 10% missingness were excluded from the analyses. 21 traits passed quality control: length, weight, abdominal fat, other fat, total fat, total cholesterol, unesterified cholesterol, free fatty acids, glucose, insulin, triglycerides, high density lipoprotein cholesterol (HDL), low density lipoprotein + very low density lipoprotein (LDL + VLDL), monocyte chemotactic protein-1 (MCP-1 / CCL2), aortic lesion size, aneurysm severity, medial aortic calcification, lateral aortic calcification, the ratio of glucose over insulin, the ratio of 100 x total fat over weight, the ratio of HDL over LDL + VLDL. Descriptions of trait measurement protocols can be found in (Ghazalpour et al., 2006), (Estrada-Smith et al., 2004), and (Meng et al., 2007).

Normalisation was achieved through a natural log transform for MCP-1, insulin, triglycerides, HDL, the ratio of glucose over insulin, and the ratio of HDL over LDL + VLDL. A square root transform was applied to the measurements of aneurysm severity, medial aortic calcification, and lateral aortic calcification due to measurements of zero for many samples.

### Network inference and module detection

Network inference and module detection were performed on a per-tissue basis using weighted gene coexpression network analysis (WGCNA) version 1.43.1 with the default parameters (Langfelder and Horvath, 2008). First, the pairwise probe coexpression was calculated for all probes passing quality control using the Pearson correlation coefficient. Next, the pairwise probe adjacencies were constructed by taking the element-wise absolute value of the coexpression and exponentiating it to the smallest power such that the connectivity of the resulting adjacency matrix was approximately scale-free (Scale-free topology criterion R^2^ > 0.85) (Zhang and Horvath, 2005). The automated selection procedure selected the exponents of 12, 5, 4, and 12 for the brain, liver, adipose, and muscle tissues respectively (**Figure S2**). Subsequently, the topological overlap dissimilarity between probe adjacencies was calculated and hierarchically clustered using the average linkage method. Hierarchically nested modules were identified from the results dendrogram using the dynamic tree cut algorithm with default parameters (Langfelder et al., 2008). Similar modules were merged together using an iterative process in which modules with summary expression profiles (eigengenes) clustering together (hierarchical average-linkage) below a height of 0.2 were joined.

Summary expression profiles were calculated as the first principal component of each module’s expression matrix. Since eigenvectors are unique up to sign, they were multiplied by the sign of their correlation with the average module gene expression within each sample (Langfelder and Horvath, 2008) to ensure eigenvector solutions were concordant with module gene expression.

### Definitions of module preservation statistics

The module preservation statistics developed by Langfelder *et al.* are defined as follows (Langfelder et al., 2011):

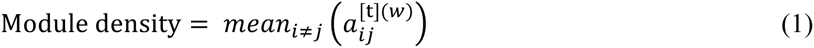

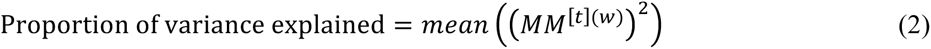

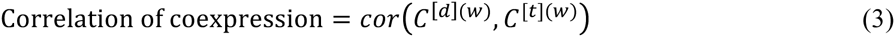

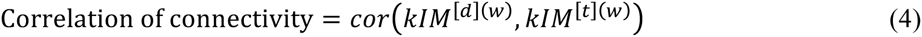

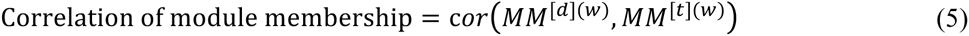

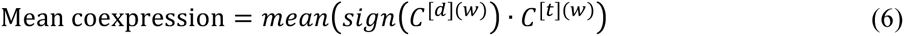

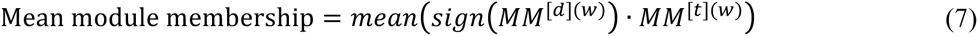

For *n* genes measured across *m* samples, *G* refers to the *m × n* matrix of gene expression, *C* refers to the *n*×*n* matrix of pairwise-gene coexpression, and *a* refers to an element of the *n × n* matrix of pairwise-gene adjacencies. The superscript *(w)* denotes the subset of genes corresponding to any given module *h* under examination and the superscripts *[d]* and *[t]* indicate whether the elements they are attached to were calculated from the *discovery* or *test* datasets respectively. The subscript letters *i* and *j* denote individual genes in the module *q*. *MM* denotes the vector of module membership for genes in module *q*, calculated as the correlation between each module gene and the module’s summary expression profile. *kIM* denotes the vector of intramodular connectivity for genes in module *q*, calculated as the sum of each module gene’s connection strength (adjacency) to all other genes in the module. The *sign* function evaluates to 1 if its argument is a positive value or −1 if its argument is a negative value.

Biological interpretations for the module preservation statistics are provided in the **Supplemental Materials.**

### Null distribution generation and permutation testing

A permutation procedure was employed to characterise each of the module preservation statistics the null hypothesis of non-preservation and to perform subsequent permutation testing. This procedure involves calculation of each module preservation statistic on *t* random gene sets of identical size to any given module under examination. Permutation test p*-*values were obtained using the estimator described in (Phipson and Smyth, 2010), which reduced to the following ratio in our analyses (see **Supplemental Materials**):

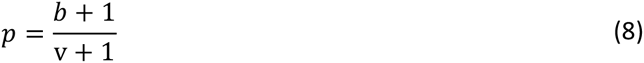

Where *v* was the total number of permutations and *b* was the number of null distribution observations more extreme than the true value for any given module preservation statistic.

For comparison with previous methodology (**Table 1**), Z-scores, summary statistic, and heuristic thresholds were calculated as previously described (Langfelder et al., 2011). Standardised Z-scores for each module preservation statistic were calculated using the mean and standard deviation estimated from the null distributions, and the combined summary score calculated as:

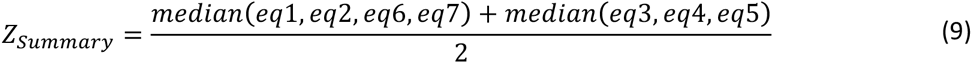

Where eq1–eq7 correspond to the module preservation statistics described by equations 1–7 above. The heuristic thresholds for significance have been defined as follows: strong evidence for module preservation if Z_summary_ > 10, weak evidence if 2 < Z_summary_ < 10, and no evidence of preservation if Z_summary_ ≤ 2 (Langfelder et al., 2011).

### Module GO term and KEGG pathway enrichment

Enrichment of Gene Ontology (GO) terms (Ashburner et al., 2000) and Kyoto Encyclopedia of Genes and Genomes (KEGG) pathways (Kanehisa and Goto, 2000) for each module were determined through overrepresentation analysis (**Tables S1 and S3**). Briefly, a hypergeometric test was performed for each GO term and KEGG Pathway annotating at least two module genes. Annotations were considered significant at the false discovery rate (FDR) corrected significance threshold of *P* < 0.05 within each annotation type (KEGG pathway, GO biological process, GO molecular function, GO cellular component).

GO term annotations for *Mus musculus* genes were obtained from the Genome-wide Annotation for Mouse Database provided through the R package *org.Mm.eg.db* (version 2.14.0). Definitions for each GO term were obtained from the GO database through the *GO.db* R package (version 2.14.0). KEGG Pathway annotations for *Mus musculus* genes were retrieved from the KEGG database using the *KEGGREST* package (version 1.4.1).

### Statistical tests

Associations between body weight and each multi-tissue module were assessed through linear regression of weight on the module’s summary expression profile adjusting for sex (**Table S2**). Multi-tissue modules are those with strong evidence for preservation in any other tissue. Associations with body weight were assessed in the module’s discovery tissue as well as any tissues for which the module had strong evidence of preservation. Effect sizes correspond to change in standard deviation (SD) of body weight per SD increase in the module’s summary expression profile in the corresponding test tissue. An association was considered significant at a Bonferroni-corrected threshold of *P* < 0.0001, adjusting for the 273 tests.

Associations between liver module 35 (LM35) and the cardiovascular disease risk traits, excluding weight, were assessed through a linear regression model adjusting for sex (**Table S4**). Summary expression profiles were calculated from adipose and muscle tissue expression for probes comprising LM35. In total 20 associations were assessed through linear regression of each trait on LM35 adipose expression, and total 20 associations were assessed through linear regression of each trait on LM35 muscle expression. Effect sizes correspond to change in LM35 tissue expression (SD-units) per SD-increase of each trait. False-discovery rate (FDR) correction was applied within each tissue to the resulting linear model p-values. We considered an association significant at FDR *q* < 0.025 (adjusting for the two tissues).

### Software and hardware

Analyses were performed using the statistical computing software, R version 3.1.3 (http://www.rproject.org/). Network module preservation was assessed using the NetRep package version 0.23.0 (https://github.com/InouyeLab/NetRep/releases/tag/v0.23.0). The latest stable version of the software can be found at http://github.com/InouyeLab/NetRep.

Analyses were performed on a Dell R910 with 40 cores (Intel Xeon 2.00 GHz), 512 Gb of RAM, 16x 1 Tb hard drives with 7200 RPM, running the 64 bit version of Ubuntu 12.0.5 LTS.

## Acknowledgements

This study was supported by funding from National Health and Medical Research Council (NHMRC) grant APP1062227. MI was supported by an NHMRC and Australian Heart Foundation Career Development Fellowship (no. 1061435). SR was supported by an Australian Postgraduate Award and a PhD student top-up award from Victorian Life Sciences Computation Initiative (VLSCI). GA was supported by an NHMRC Early Career Fellowship (no. 1090462).

The Mouse Model of sexually Dimorphic Atherosclerotic traits data was generated and was contributed by Jake Lusis, Eric Schadt and Merck Pharmaceutical through the Sage Bionetworks Repository.

